# Rules of tissue packing involving different cell types: human muscle organization

**DOI:** 10.1101/038968

**Authors:** Daniel Sánchez-Gutiérrez, Aurora Sáez, Carmen Paradas, Luis M. Escudero

**Affiliations:** Departamento de Biología Celular, Universidad de Sevilla and Instituto de Biomedicina de Sevilla (IBiS), Hospital Universitario Virgen del Rocío/CSIC/Universdad de Sevilla. 41013 Seville, Spain.; Dpto. Teoría de la Señal y Comunicaciones. Universidad de Sevilla. Cmno. de los descubrimientos s/n 41092, Sevilla; Instituto de Biomedicina de Sevilla (IBiS), Hospital Universitario Virgen del Rocío/CSIC/Universdad de Sevilla. 41013 Seville, Spain.

## Abstract

Natural packed tissues are assembled as tessellations of polygonal cells that do not leave empty spaces between them. They include the epithelial sheets and the skeletal muscles. Epithelia are formed by equivalent cells that change shape and organization through development. The skeletal muscles appear as a mosaic composed by two different types of cells: the slow and fast fibres that are determined by the identities of the motor neurons that innervate them. Their relative distribution is important for the muscle function and can be altered in some neuromuscular diseases. Little is known about how the spatial organization of fast and slow fibres is established and maintained. In this work we use computerized image analysis and mathematical concepts to capture the organizational pattern in two different healthy muscles: biceps brachii and quadriceps. Here we show that each type of muscle portrays a characteristic topological pattern that allows distinguishing between them. The biceps brachii muscle presents a particular arrange based on the different size of slow and fast fibres, contrary to the quadriceps muscle where an unbiased distribution exists. Our results indicate that the relative size of each cellular type imposes an intrinsic organization into the tissue. These findings establish a new framework for the analysis of packed tissues where two or more cell types exist.

## INTRODUCTION

The organization of the cells in the tissue is perfectly controlled to drive major shape changes during morphogenesis. All these events lead to the final arrangement of cells what it is closely related to organ function. Packed tissues have been used as a model to understand the key processes on the establishment of the function of an organ (Chichilnisky, 1986; Classen et al., 2005; Gibson et al., 2006; Hayashi and Carthew, 2004; Honda, 1978; Korn and Spalding, 1973; Lewis, 1928; Pilot and Lecuit, 2005; Rivier et al., 1995). Most of these works have been based in the idea that the apical regions of the epithelial cells were polygons and the epithelial organization was analyzed using the distribution of sides of their cells. Skeletal muscles are also packed tissues composed by closely arranged fibres separated by a fine layer of conjunctive tissue (the endomisium) (Helliwell, 1999). A muscle biopsy section appears as a mosaic of fibres that also organize as polygons in a tessellation (leaving no empty space between them). Therefore they also can be analyzed in terms of organization as epithelial tissues (Sáez et al., 2013b; Sanchez-Gutierrez et al., 2016).

We have recently introduced network theory and Centroidal Voronoi tessellations (CVT) to the study of packed tissue organization (Escudero et al., 2011; Sanchez-Gutierrez et al., 2016). In these reports, mathematical concepts were used to objectively quantify the organization of natural packed tissues such epithelia or muscles. The comparison with the CVT added a new insight since it was possible to infer some biophysical properties from the packed tissues that were also supported by computer simulations. Packed tissues obey several laws that related area and organization. These includes the Euler’s Theorem that estates that the average number of neighbours of a cell will be close to six; the Lewis’ law that linearly relates the average area of a cell with its number of sides (so, small cells tend to have less sides, and big cells tend to have higher number of sides); and the Aboav-Weaire law that establish an inverse relationship between the average number of sides of a cell and the average number of sides of their neighbours (Aboav, 1970; Chiu, 1995; Gervois et al., 1992; Hilhorst, 2006; Lewis, 1928; Rivier et al., 1995; Weaire, 1974). In addition, it was shown that there is a physical constrain affecting natural packed tissues that restrict them to certain organizations. These arrangements are similar to the polygons distributions that present the CVT.

All the previous studies investigating tissue organization have considered tissues formed by cells with the same properties and capabilities: equivalent entities that could transiently vary the properties depending of the cell cycle stage or changes in the cytoskeleton (Aegerter-Wilmsen et al., 2012; Farhadifar et al., 2007; Levayer and Lecuit, 2013; Mao et al., 2013; Sanchez-Gutierrez et al., 2016; Zallen and Zallen, 2004). Here we analyze the organization of skeletal muscle tissues considering the distribution of myofibres into fast and slow twitch type (Pette and Staron, 2000) which are determined by the specific myosin protein expressed in each fibre. This establishes a mosaic or “checked” pattern that is a characteristic feature of skeletal muscle. Although some muscles have a higher proportion of one of these types of fibres, most of them would be almost indistinguishable one from another based on the proportions of the type of fibres.

The neuromuscular system is constituted by the muscle-controlling neurons in the spinal cord, the peripheral motor neurons, the neuromuscular junctions and the muscles themselves. Neuromuscular diseases are a large group of pathologies produced by the affection of one or more of these components, with very heterogeneous etiology and course. The evaluation of the changes in the morphological characteristics of a given biopsy with respect the normal muscle is one of the main features for the diagnostic of a neuromuscular disorder (Clarke and North, 2003; Dubowitz, 1974; Dubowitz and Sewry, 2007; Na et al., 2006). Morphological pathogenic features to evaluate in a muscle biopsy include alterations of fibre size, nuclei position, and amount of connective tissue or necrotic fibres. Changes of the distribution pattern of slow and fast fibres can also be detected, being a typical feature of the neurogenic disorders such neuropathies or amyotrophic lateral sclerosis (Dubowitz and Sewry, 2007; Sáez et al., 2013b). In addition, switch from fast to slow twitch type fibre and predominance of one fibre type, or even uniformity of fibre types, are detected in some types of myopathies (De Palma et al., 2006).

Since the way the skeletal muscle degenerates under pathogenic conditions is critical to determine the cause of many neuromuscular disorders, the accurate definition of the features in normal muscles is also essential to better identify the disease. Considering that most of muscle biopsies are taken from biceps brachii and deltoids muscles in upper limbs, and quadriceps, tibialis anterior and gastrocnemius in lower limbs these are the muscles that should be described under normal conditions from a clinical perspective. To analyse the structural and organizational pattern of skeletal muscles, a high amount of samples is mandatory (Escudero et al., 2011; Sáez et al., 2013a; Sáez et al., 2013b). Therefore, we selected biceps brachii and quadriceps muscles because the amount of available normal samples and the morphological similarity between them regarding distribution of the type of fibre.

In this work we integrate geometric and topological data to capture an organizational signature in packed tissues with two different cellular types. Our results indicate that biceps brachii and quadriceps can be distinguished based the pattern of slow and fast cells. We demonstrate that the mosaic these two cell types defines a differential organization for skeletal muscles.

## RESULTS

### Computerized analysis of biceps brachii and quadriceps biopsy images

We have compared biceps brachii (BA) and quadriceps (QA) muscles from control male adult individuals in terms of morphological characteristics of their fibres. Thin sections of biopsies were analyzed using immunohistochemical staining. We combined anti-collagen VI antibody that provides the outline of the muscle fibres (and enables the quantification of the amount of collagen in the tissue) and, anti-myosin slow (type I) or anti-myosin fast (type II) specific antibodies that allow the identification of fibre type **(Fig. 1).** In the case of BA, 18 biopsies were analyzed, obtaining 34 micrographs and 90 Region Of Interest (ROI) **(Fig. 1 A-C** and **Table S1).** 6 QA biopsies were used to obtain 9 micrographs and 25 ROI **(Fig. 1 D-F** and **Table S1).** The human visual analysis of the different ROI makes very difficult to extract patterns that could differentiate both types of muscles **(Fig. 1**). We used a computerized approach aiming to capture a characteristic signature form each one of them. First, the images were segmented to identify the outline of the fibres and the content of collagen (Sáez et al., 2013a; Sáez et al., 2013b). Then the values for a series of 14 geometrical characteristics (14 first features in **Table 1**) and the proportion of slow cells (feature 69 in **Table 1**) were calculated. In each type of muscle, samples were very heterogeneous and presented a wide range of values for each characteristic. We started examining some of these geometric characteristics and comparing their averages values between both types of muscles **(Table S2).** BA fibres were approximately 33% bigger than QA in average. Both BA and QA presented a lower proportion of slow fibres (around the 31% and 25% respectively) that fast fibres. Interestingly, in the case of QA, the average area of fast and slow fibres was virtually the same; meanwhile in the case of BA the average size was bigger in the fast fibres compared with the slow fibres **(Table S2).**

**Table 1.** List of characteristics analyzed in this study. Table shows the name of the 69 characteristics analyzed in the study. These characteristics can be classified into three types: geometrical characteristics, related to the size and shape of cells (1-14), network characteristics, capturing the organization of the cells (15-68) and the proportion of slow cells (69). The characteristics labelled in bold are the 35 features related to the fast or slow cell type. S. D. = Standard Deviation.

| CHARACTERISTICS |  |  |  |
| --- | --- | --- | --- |
| cc | Name | cc | Name |
| 1 | Average Area | 35 | Average Strengths |
| 2 | S. D. Area | 36 | S. D. Strengths |
| 3 | **Average Area of slow cells** | 37 | **Average Strengths of fast cells** |
| 4 | **S. D. Area of slow cells** | 38 | **S. D. Strengths of fast cells** |
| 5 | **Average Area of fast cells** | 39 | **Average Strengths of slow cells** |
| 6 | **S. D. Area of fast cells** | 40 | **S. D. Strengths of slow cells** |
| 7 | Average major Axis | 41 | Average Clustering Coefficient |
| 8 | Average minor Axis | 42 | S. D. Clustering Coefficient |
| 9 | Average Relation Axis | 43 | **Average Clustering Coefficient of fast cells** |
| 10 | S. D. Relation Axis | 44 | **S. D. Clustering Coefficient of fast cells** |
| 11 | Average Convex Hull | 45 | **Average Clustering Coefficient of slow cells** |
| 12 | S. D. Convex Hull | 46 | **S. D. Clustering Coefficient of slow cells** |
| 13 | Average Relation A1/A2 | 47 | Average Eccentricity |
| 14 | S. D. Relation A1/A2 | 48 | S. D. Eccentricity |
| 15 | Average Neighbours | 49 | **Average Eccentricity of fast cells** |
| 16 | S. D. Neighbours | 50 | **S. D. Eccentricity of fast cells** |
| 17 | **S. D. Neighbours of slow cells** | 51 | **Average Eccentricity of slow cells** |
| 18 | **S. D. Neighbours of fast cells** | 52 | **S. D. Eccentricity of slow cells** |
| 19 | **slow Neighbours of slow cells** | 53 | Average Betweenness Centrality |
| 20 | **fast Neighbours of slow cells** | 54 | S. D. Betweenness Centrality |
| 21 | **slow Neighbours of fast cells** | 55 | **Average Betweenness Centrality of fast cells** |
| 22 | **fast Neighbours of fast cells** | 56 | **S. D. Betweenness Centrality of fast cells** |
| 23 | Average Relation Neighbours Area | 57 | **Average Betweenness Centrality of slow cells** |
| 24 | S. D. Relation Neighbours Area | 58 | **S. D. Betweenness Centrality of slow cells** |
| 25 | Average Relation Neighbours major axis | 59 | Average Shortest Paths lengths |
| 26 | S. D. Relation Neighbours major axis | 60 | S. D. Shortest Paths Lengths |
| 27 | Average Relation Neighbours minor axis | 61 | **Average Shortest Paths Lengths from fast cells to fast cells** |
| 28 | S. D. Relation Neighbours minor axis | 62 | **S. D. Shortest Paths Lengths from fast cells to fast cells** |
| 29 | Average Relation Neighbours relation axis | 63 | **Average Shortest Paths Lengths from fast cells to slow cells** |
| 30 | S. D. Relation Neighbours relation axis | 64 | **S. D. Shortest Paths Lengths from fast cells to slow cells** |
| 31 | Average Relation Neighbours convex hull | 65 | **Average Shortest Paths Lengths from slow cells to slow cells** |
| 32 | S. D. Relation Neighbours convex hull | 66 | **S. D. Shortest Paths Lengths from slow cells to slow cells** |
| 33 | Average Relation Neighbours relation A1/A2 | 67 | **Average Shortest Paths Lengths from slow cells to fast cells** |
| 34 | S. D. Relation Neighbours relation A1/A2 | 68 | **S. D. Shortest Paths Lengths from slow cells to fast cells** |
|  |  | 69 | **Proportion of slow cells** |

**Figure 1.**
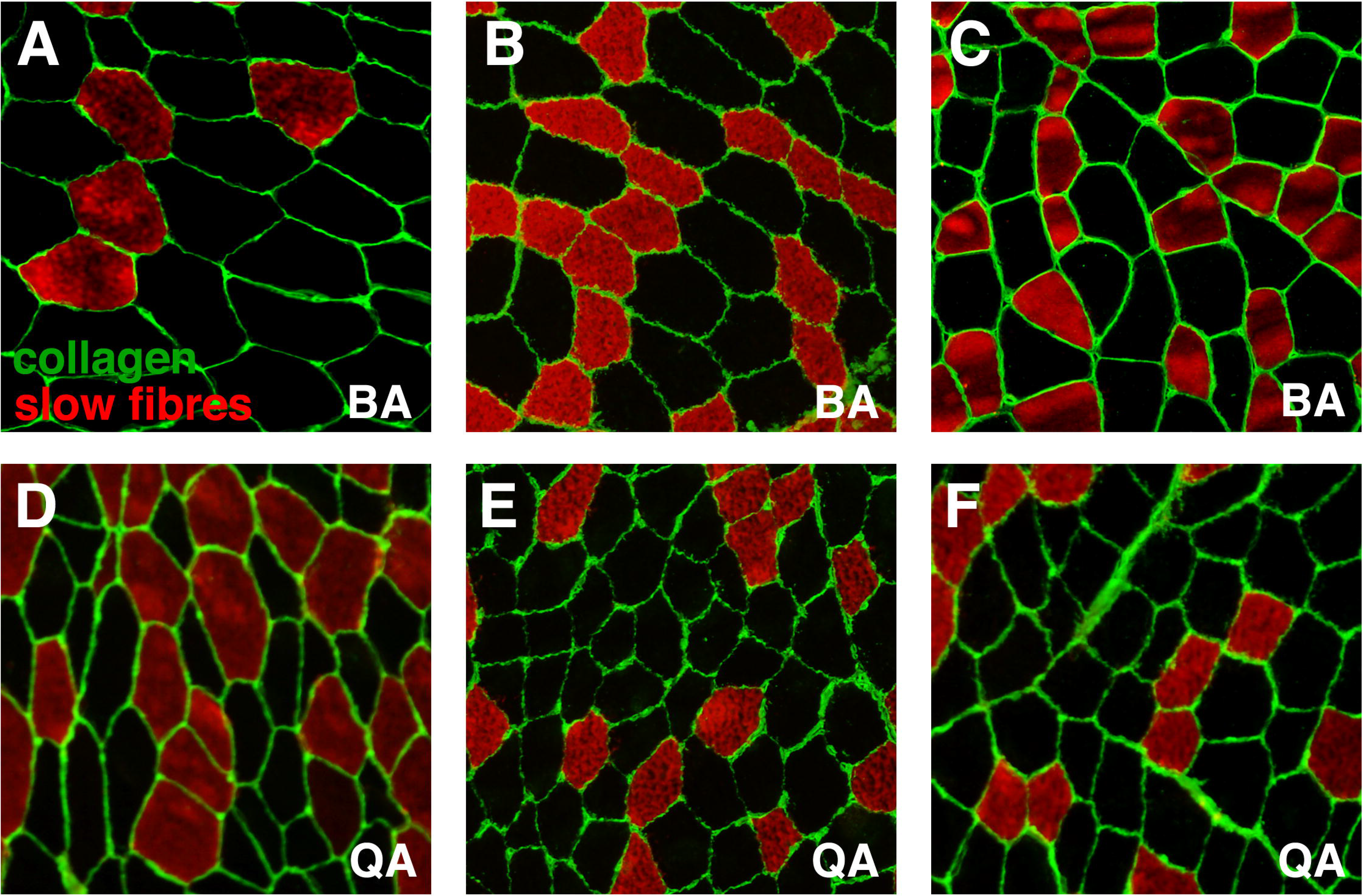
images from control human muscle biopsies. Fluorescence images corresponding to control biopsies showing collagen VI content including the endomysium and perimysium (green), slow fibres (red) and fast fibres (black). **(A, B, C)** Images from control biceps brachii. **(E, F, G)** Images from control quadriceps.

### Biceps brachii and Quadriceps present different organization of fibres

We compared the organization of BA and QA samples using a network approach that evaluates topological characteristics aiming to identify small organizational differences between similar images (Sáez et al., 2013b; Sanchez-Gutierrez et al., 2013). The method is based in the consideration of the tissue as a network of cell to cell contacts (Escudero et al., 2011). Under this premise, we extracted the values for 54 “network” characteristics (features 15-68 in **Table 1)** besides the 14 geometric features and the proportion of slow cells. In this way, we obtained a vector of 69 features for each muscle ROI. Due to the large difference in the number of ROIs (90 BA vs. 25 QA) we designed a protocol to use the whole data and at the same time be able to obtain comparable results. The protocol consisted in performing 1,000 combinations of ROIs. Each combination was done using 25 images of each group. To obtain a baseline for our evaluation system, we first performed 1,000 comparisons using only BA images: Two groups of 25 BA images were chosen randomly from the total 90 each time. Each comparison was used to perform a feature selection step that chose the most relevant characteristics from the total of features assayed. The selected features were used to perform a Principal Component Analysis (PCA) and obtain a value for the “PCA descriptor” that quantified the degree of separation between both groups of images (Sanchez-Gutierrez et al., 2013) and **Materials and Methods).** The values of the PCA descriptor ranged from 0.08 to 0.88, and presented a median value of 0.23 **(Fig. 2A).** We then performed other 1,000 randomizations: in each one, 25 images from the 90 BA were selected and compared with the 25 QA images. In this case the values of the PCA descriptor ranged from 0.49 to 2.79, with a median value of 1.09 **(Fig. 2B).** Comparing the PCA graphs corresponding to the median and best values, the separation was largely improved in the case of BA-QA with respect BA-BA **(Fig. 2B** and **Fig. 2A).** We also observed that the BA-QA values were lower when using only the 15 geometric characteristics (ranging from 0.25 to 2.25, with a median value of 0.76, **(Fig. 2C)**, indicating the importance of the network characteristics to improve the separation.

**Figure 2.**
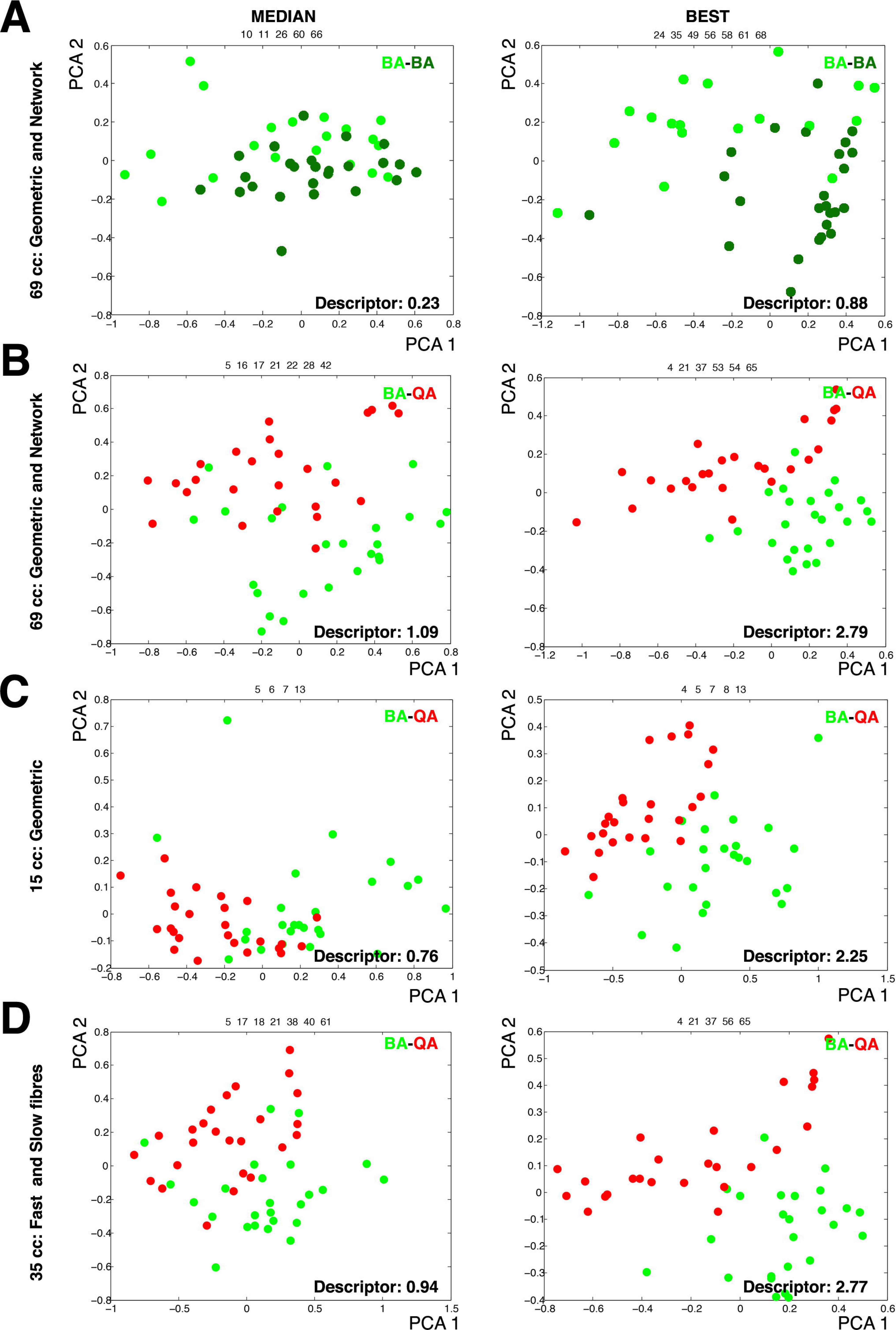
Principal component analysis graphs for different combinations of muscle type images and characteristics. PCA graphs for the comparisons two groups of 25 images. Selected panels were the combination that presents the median (left) and best (right) descriptor using different sets of characteristics. The green dots (dark or light) represent BA images. The red dots represent QA images. **A)** 25 images randomly taken from a set of 90 samples of BA vs other different 25 images using the set of 69 cc. **B)** 25 images randomly taken from a set of 90 samples of BA versus 25 QA images using the set of 69 cc. **C)** 25 images randomly taken from a set of 90 samples of BA versus QA images using the set of 15 cc. **D)** 25 images randomly taken from a set of 90 samples of BA versus QA images using the set of 35 cc.

### Similar muscles differ in the organization of fast and slow fibres

We examined the features that were relevant to separate BA and QA samples trying to understand the biological differences between these two similar muscles. Each feature selection step selects a maximum of 7 features per comparison. We calculated the rate of appearance of each feature in each one of the 1,000 comparisons performed in each case. In our baseline assay, the 1,000 BA-BA comparisons, we did not find clear predominant characteristics being the maximum frequency a 20.6% of the randomizations **(Table 2**, features above the 15% of frequency). We compared these results with the BA-QA assay. In this case there was a clear predominance of some characteristics over others **(Table 2**, features above the 25% of frequency). This indicated that different combinations of BA images could be separated from QA images using the same features. In short, these results suggest the existence of some general differences between BA and QA. The most frequent characteristics appearing in the BA-QA comparisons were mainly related to the geometry or organization of the types of fibres (the nine most frequent features in **Table S3).** In particular, the “standard deviation of the area of the slow cells” and the “number of slow neighbours of fast cells” were the two more relevant features. This suggested that the difference between BA and QA could stem on the distribution of fast and slow fibres. To test this idea, we repeated the 1,000 BA-QA comparisons using only the 35 characteristics that were specifically related to fast and slow fibres. The distribution of values for the PCA descriptor was still high (ranging from 0.35 to 2.77, with a median value of 0.94, **Fig. 2D).** We also observed the predominance of the same type of features than in the experiment with 69 characteristics **(Table S3).**

**Table 2.** Frequency of characteristics that better differentiate BA and QA images. This table shows the characteristics that have been selected with a higher frequency in the 1,000 BA-BA and BA-QA comparisons (using 69 characteristics).

| **Feature selection: 1,000 combinations from BA (25 random images) vs BA (25 random images)** |  |  |
| --- | --- | --- |
| **Name** | **Characteristic** | **Frequency** |
| S. D. Shortest Paths Lengths from fast cells to slow cells | 64 | 20.60% |
| Average Shortest Paths Lengths from slow cells to slow cells | 65 | 19.60% |
| Average Neighbours | 15 | 16.60% |
| S. D. Shortest Paths Lengths from slow cells to fast cells | 68 | 15.30% |
| S. D. Betweenness Centrality of slow cells | 58 | 15% |

| **Feature selection: 1,000 combinations from BA (25 random images) vs QA** |  |  |
| --- | --- | --- |
| **Name** | **Characteristic** | **Frequency** |
| S. D. Area of slow cells | 4 | 67.20% |
| Slow Neighbours of fast cells | 21 | 48.40% |
| Average Strengths of fast cells | 37 | 40.20% |
| Average Strengths of slow cells | 39 | 36.80% |
| Average Area of fast cells | 5 | 33.80% |
| S. D. Neighbours of slow cells | 17 | 28.40% |

### Importance of the proportion of fast and slow cells on the muscle organization

We have shown that BA and QA present different average proportions of fast and slow fibres **(Table S2).** This influence the values of the network characteristics related to the slow and fast fibres. We tried to evaluate the importance of these proportion differences in the muscle organization. To do that, we selected two groups of 25 BA images with very different percentage of slow fibres. Using 69 characteristics for the comparison the PCA graph showed two clearly separated groups, and the PCA descriptor value was extremely high: 11.33 **(Figure 3A).** In this case, the difference between the average percentages of slow cells between these two groups was 0.216 (we will call this value Δ proportion). In parallel, we compared QA samples with a selection of BA samples with the percentage of slow cells more similar to QA (a Δ proportion value of 0.002). In this case there was some degree of separation with a descriptor of 1.07 when using 69 characteristics **(Figure 3B, Table 3).** Interestingly, this value was very similar to the median value of the 1,000 BA-QA comparisons (1.09; **Fig. 2B).** To further investigate the relation between Δ proportion and the separation of the groups of images we took 1,000 BA-BA comparisons to plot the values for the PCA descriptor against its correspondent Δ proportion values **(Figure 3C).** We observed a poor relation between the increase of the Δ proportion and the PCA descriptor (Pearson’s coefficient r= 0.2435). The same happened when we used the 1,000 BA-QA comparisons **(Figure 3D**, Pearson’s coefficient r= 0.2735). These results suggested that the proportion of slow cells is not the main factor responsible for the differences between BA and QA tissues.

**Table 3.**

| Biceps brachii Adult |  | Original | Random slow/fast cells |  |  |
| --- | --- | --- | --- | --- | --- |
| Number | Characteristic | Value | Min | Max | Median |
| 3 | Average Area of slow cells | 23616.36 | 25488.99 | 26776.38 | 26080.13 |
| 4 | S. D. Area of slow cells | 4242.18 | 5951.76 | 6964.85 | 6403.99 |
| 5 | Average Area of fast cells | 27392.63 | 25643.48 | 26536.55 | 26076.49 |
| 6 | S. D. Area of fast cells | 6409.93 | 6167.93 | 6785.72 | 6495.91 |
| 17 | S. D. Neighbours of slow cells | 0.78 | 0.79 | 0.90 | 0.84 |
| 18 | S. D. Neighbours of fast cells | 0.85 | 0.80 | 0.89 | 0.85 |
| 19 | slow Neighbours of slow cells | 1.65 | 2.56 | 2.82 | 2.69 |
| 20 | fast Neighbours of slow cells | 4.26 | 3.19 | 3.43 | 3.32 |
| 21 | slow Neighbours of fast cells | 1.95 | 2.64 | 2.84 | 2.74 |
| 22 | fast Neighbours of fast cells | 4.14 | 3.15 | 3.36 | 3.26 |
| 37 | Average Strengths of fast cells | 1197.19 | 1152.29 | 1183.17 | 1167.90 |
| 38 | S. D. Strengths of fast cells | 249.06 | 235.00 | 261.19 | 247.81 |
| 39 | Average Strengths of slow cells | 1123.39 | 1145.36 | 1189.76 | 1167.80 |
| 40 | S. D. Strengths of slow cells | 221.58 | 228.91 | 266.04 | 245.73 |
| 43 | Average Clustering Coefficient of fast cells | 69.49 | 69.70 | 71.68 | 70.73 |
| 44 | S. D. Clustering Coefficient of fast cells | 17.55 | 16.66 | 18.48 | 17.62 |
| 45 | Average Clustering Coefficient of slow cells | 72.27 | 69.13 | 72.45 | 70.73 |
| 46 | S. D. Clustering Coefficient of slow cells | 16.38 | 16.11 | 19.03 | 17.43 |
| 49 | Average Eccentricity of fast cells | 2102.00 | 2079.41 | 2112.54 | 2095.04 |
| 50 | S. D. Eccentricity of fast cells | 269.37 | 254.36 | 276.71 | 264.78 |
| 51 | Average Eccentricity of slow cells | 2080.82 | 2074.78 | 2117.68 | 2095.31 |
| 52 | S. D. Eccentricity of slow cells | 250.47 | 248.99 | 277.91 | 263.52 |
| 55 | Average Betweenness Centrality of fast cells | 1324.80 | 1221.65 | 1438.54 | 1314.93 |
| 56 | S. D. Betweenness Centrality of fast cells | 1004.46 | 791.91 | 1195.70 | 965.58 |
| 57 | Average Betweenness Centrality of slow cells | 1330.29 | 1227.92 | 1415.92 | 1316.04 |
| 58 | S. D. Betweenness Centrality of slow cells | 971.95 | 801.17 | 1140.23 | 978.55 |
| 61 | Average Shortest Paths Lengths from fast cells to fast cells | 683.97 | 662.75 | 693.15 | 676.53 |
| 62 | S. D. Shortest Paths Lengths from fast cells to fast cells | 309.38 | 299.37 | 312.65 | 305.66 |
| 63 | Average Shortest Paths Lengths from fast cells to slow cells | 789.92 | 707.30 | 772.82 | 739.85 |
| 64 | S. D. Shortest Paths Lengths from fast cells to slow cells | 273.72 | 277.99 | 301.80 | 288.77 |
| 65 | Average Shortest Paths Lengths from slow cells to slow cells | 663.20 | 653.53 | 697.28 | 676.70 |
| 66 | S. D. Shortest Paths Lengths from slow cells to slow cells | 290.31 | 291.45 | 313.76 | 300.95 |
| 67 | Average Shortest Paths Lengths from slow cells to fast cells | 645.91 | 682.38 | 724.70 | 703.57 |
| 68 | S. D. Shortest Paths Lengths from slow cells to fast cells | 301.25 | 285.64 | 302.72 | 294.66 |

Comparison of real values and random values for each characteristic and type of muscle. The table shows results of the evaluation of the 34 characteristics related with the fast and slow condition of the fibres. Each original value is compared with the minimum, maximum and median values for 10,000 randomizations. The original values labelled in bold mark the ones outside of the range of values of the random distribution in each case.

**Figure 3.**
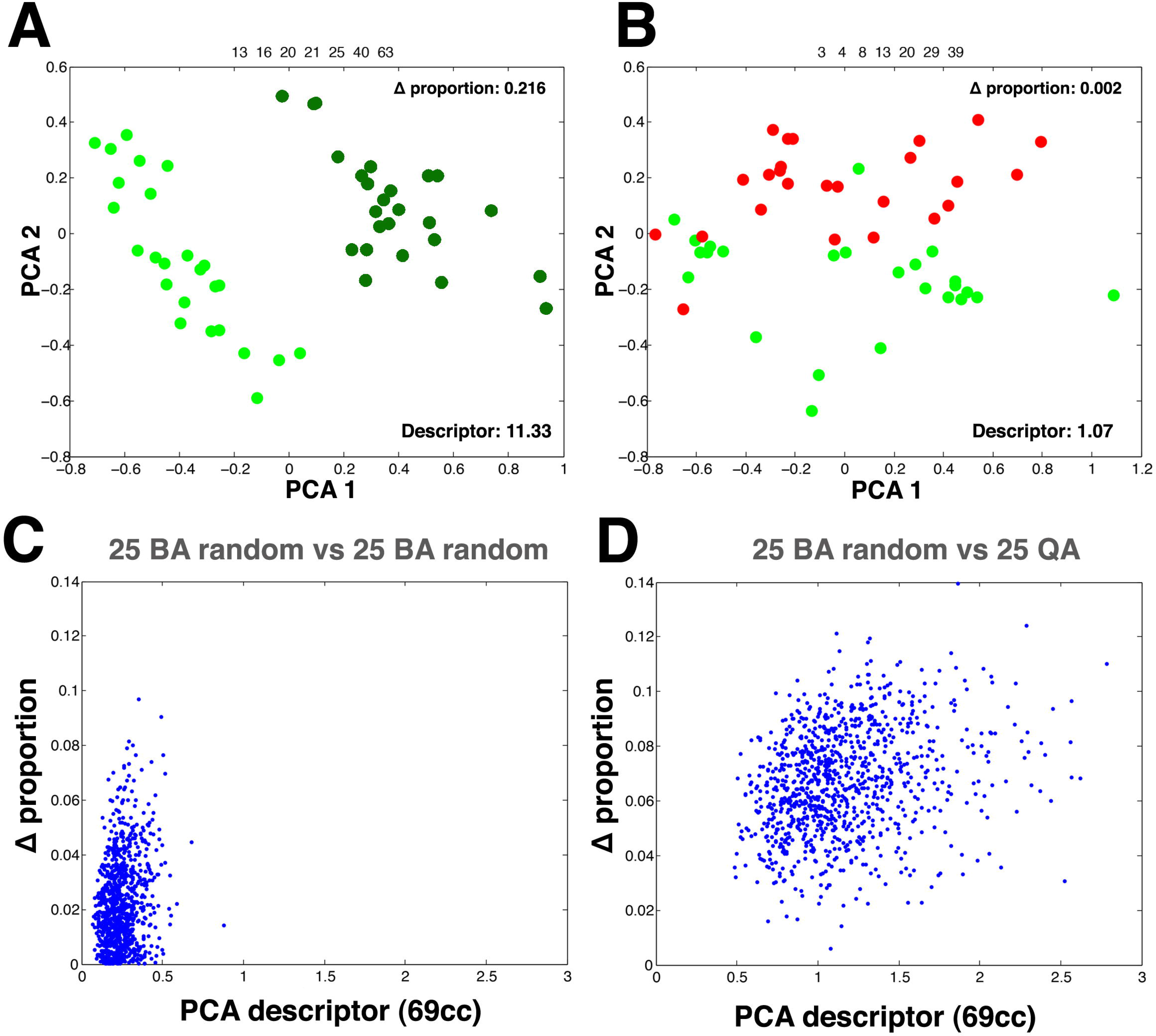
Influence of the proportion of slow fibres in the muscle organization. A) Comparison of 25 images from BA (light green dots) vs 25 images from BA (dark green dots) using two groups of BA images with a very different percentage of slow fibres (Δ proportion= 0.216) and a set of 69 characteristics. The result is a clear separation of both groups with a PCA descriptor of 11.33. **B)** Comparison of 25 images from BA (green dots) with very similar percentage of slow fibres (Δ proportion= 0.002) than the 25 QA images (red dots) and a set of 69 characteristics. The graph shows some overlap between the two groups (PCA descriptors= 1.07). **C)** Graph representing the 1,000 random comparison of 25 images random from BA versus 25 images random from BA (blue dots). “Δ proportion” of slow fibres is represented against the PCA descriptor value of the same random comparison. **D)** Graph representing the 1,000 random comparison of 25 images random from BA versus QA (blue dots). “Δ proportion” of slow fibres is represented against the PCA descriptor value of the same random combination.

### The relative size of slow and fast fibres affects their relative distribution

The muscles fibres are arranged in bundles and the sections of the muscular tissue analysed are similar to tessellations of convex polygons. This property has been previously used to try to capture the organization of packed tissues (Classen et al., 2005; Farhadifar et al., 2007; Gibson and Gibson, 2009; Sanchez-Gutierrez et al., 2016). We examined our biceps brachii and quadriceps samples in these terms and found that they presented a similar polygon distribution **(Fig. 4A** and **Table S4;** MANOVA *p value* = 0.3196). In packed cellular arrangements the area and the number of neighbours are related by several general rules such as Euler’s theorem and Lewis and Aboav-Weaire laws (Aboav, 1970; Chiu, 1995; Gervois et al., 1992; Hilhorst, 2006; Lewis, 1928; Rivier et al., 1995; Weaire, 1974). As we have mentioned before the one of the clearest differences between BA and QA samples was the average relative size between fast and slow fibres. We examined if this difference was extended to the distribution of fibres size **(Fig. 4B).** In the case of QA both distributions presented a very high level of overlapping (**Fig. 4B** left panel). In the other hand, BA distributions of slow and fast cell areas were slightly displaced since a considerable part of the population of slow cells was smaller than the fast cells **(Fig. 4B** right panel). Although in both cases we were not able to find significant differences between slow and fast fibres area distribution (Kolmogorov-Smirnov test; QA: *p value* =1; BA: *p value* = 0.3309) we decided to continue the analysis on the relation between area distribution and organization. Following the principles of the Lewis and Aboav-Weaire laws the small difference in area distribution of slow and fast cells in BA could bias their organization: bigger cells (fast) should tend to have a higher number of neighbours, and these neighbours should tend to be smaller cells with a lower number of sides (slow). Therefore, we analyzed the polygon distribution of both types of fibres in the QA and BA images **(Fig. 4C, D** and **Table S4).** Using MANOVA test to compare slow and fast polygon distributions we were not able to find significant differences in the case of QA **(Fig 4C**, MANOVA *p value* = 0.1434). On the contrary, BA samples presented distributions significantly different **(Fig 4D**, MANOVA *p value* = 0.0037). In addition, we statistically compared the frequency of each polygon class between slow and fast fibres **(Materials and Methods).** Again, there were not differences in the case of QA **(Table S4).** In BA, we found that the amount of slow fibres that were heptagons and octagons was significantly lower than among fast fibres **(Fig. 4D** and **Table S4).** We think that the small differences in the area distribution founded in the BA samples imposed a degree of order in the BA organization that it is absent in QA.

**Figure 4.**
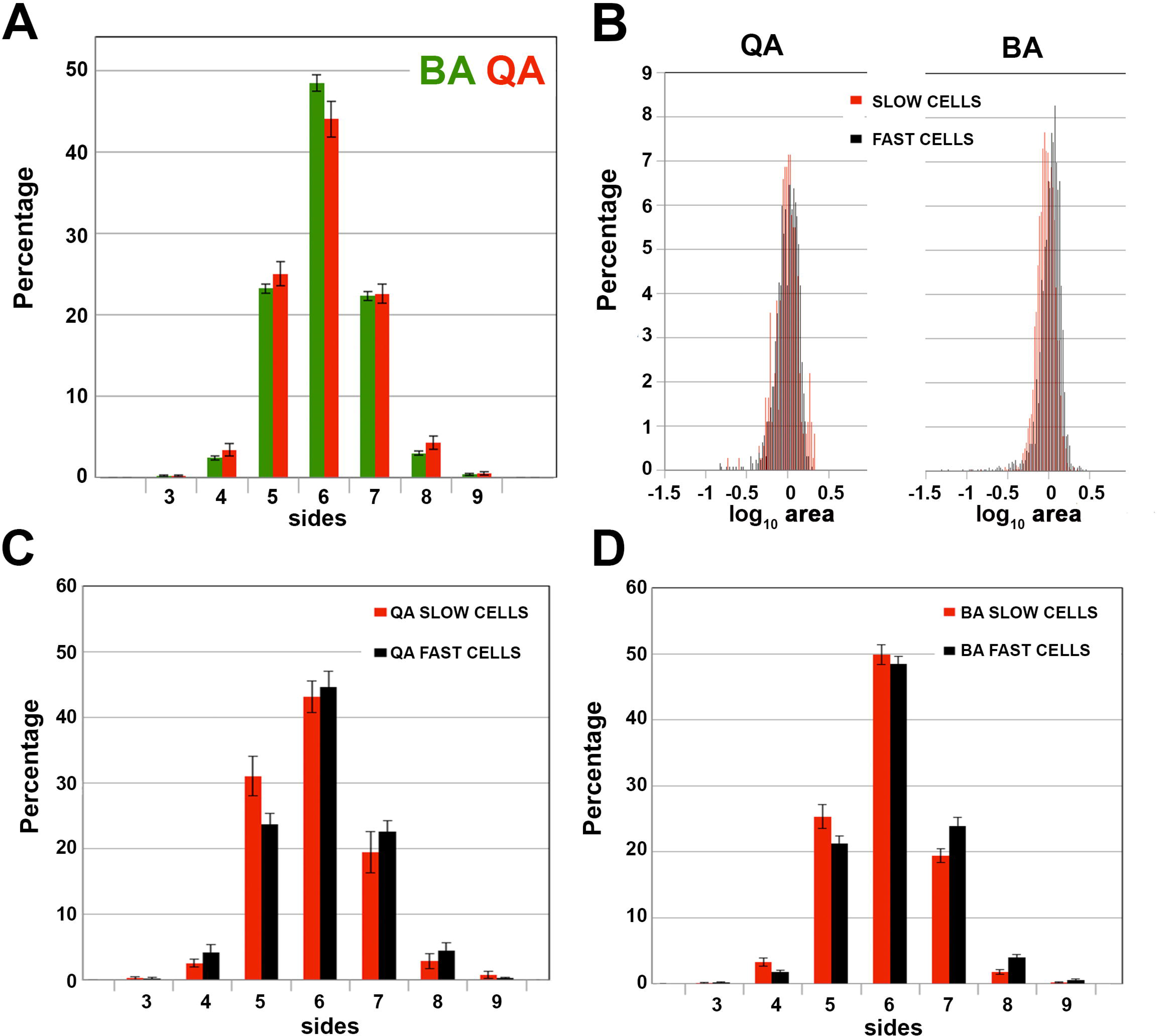
BA and QA present differences in polygon and area distribution of their slow and fast fibres. Polygon distribution of BA fast fibres (black) and BA slow fibres (red). The error bars represent the standard error. **B)** Comparison of the area distribution of QA fast and slow cells (left panel) and the area distribution of BA fast and slow cells (right panel). **C)** Polygon distribution of QA fast and slow fibres. **D)** Polygon distribution of BA fast and slow fibres. The frequency of each type of polygons in both sets of images is represented. The error bars represent the standard error.

### Slow and fast fibres present an intrinsic organization in the biceps brachii

Our diverse results suggested that BA and QA samples presented differences related to the organization of their two types of fibres. To test this hypothesis we performed simulations where in each ROI, every cell became fast or slow randomly (maintaining the percentage of fast and slow fibres constant). Logically, this changed the values for the 34 characteristics specifically related to fast and slow fibres properties. We obtained the average value for each characteristic putting together all the images of each category (90 ROI in the case of BA and 25 for QA). Then we plotted the distribution of the values for each characteristic and compared them with the distribution of values for 10,000 randomizations of fibre type **(Fig. 5A-F** and **Table 3).** We expected that if a characteristic was not affected by the fibre-type randomization, the real value would be falling inside of the distribution of random values. This was the case for all except two of the characteristics when analyzing QA samples **(Fig. 5A-C** and **Table 3).** In contrast, more than a half of BA characteristics presented the real value displaced from the distribution of random data **(Fig. 5D-E** and **Table 3).** In some cases, the real value was very different from the randomized. For example, the real average number of “slow neighbours of slow cells” was clearly lower than any of the randomized **(Fig. 5D).** This suggested that slow cells in the BA muscle were mainly surrounded by fast cells and not by other slow cells (i.e. slow fibres tended to appear isolated and the randomization grouped them). This result supported that BA organization of fast and slow fibres was not arbitrary.

**Figure 5.**
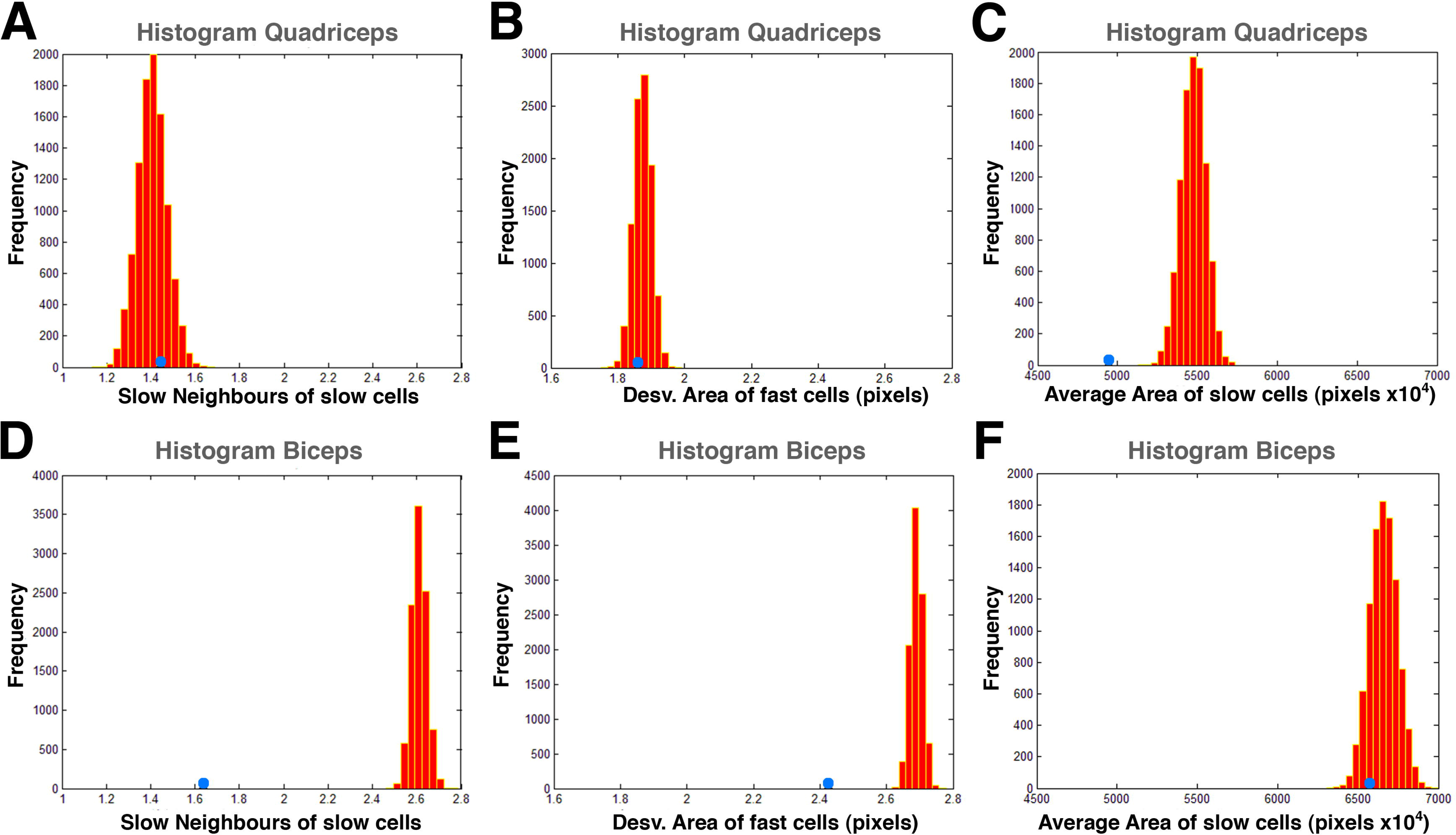
Frequency of values for characteristics depending of the distribution of fast and slow fibres. The histograms show the frequency of values for several characteristics related to the distribution of fibres type from 10,000 randomizations of the fast and slow fibres. Blue circle shows the value of the characteristic for the real distribution of fast and slow cells in the muscle. **A)** and **D)** Histogram for the characteristic “slow neighbours of slow cells” in QA and BA respectively. The real value is similar to the median of random values. **B)** and **E)** Histogram for the characteristic “deviation area of fast cells in QA and BA respectively. **C)** and **F)** Histogram for the characteristic “average area of slow cells” in QA and BA respectively. In the cases **C, D** and **E** the real value is lower than the random values.

## DISCUSSION

### Biceps brachii and quadriceps are different in terms of the organization of slow and fast fibres

In this study we integrate and quantify information from two large sets of images from two healthy muscles. Although both sets of images present high similarity after visual inspection **(Fig. 1)**, the first problem that we have encountered is the wide heterogeneity between samples from the same type of muscle. For example, the “average area” of quadriceps fibres is bigger than the “average area” of biceps brachii fibres, but at the same time a high proportion of biceps brachii images present fibres bigger than quadriceps fibres **(Table S1** and **Table S2).** In this work, we have tackle this problem using several approaches that try to incorporate all the useful data from the two sets of images. The first step has been to design a protocol to evaluate all the images available (90 BA and 25 QA), in this way for each comparison of BA and QA data we obtain 1,000 values for the PCA descriptor, and we are able to analyze the differences or similarities among all the samples. Our first conclusion is that our method is not able to completely separate both types of images. The representative graphs of the median values of the PCA descriptor show how some BA images are very similar to the QA **(Fig. 2B** and **2D** right panels). Even the best combinations still present some overlapping of images in the PCA graph **(Fig. 2B** and **2D** left panels). However we have been able to extract some useful information from these assays: i) topological characteristics improve the separation of the two groups; ii) the characteristics related to the fast and slow fibres contain most of the relevant information to better distinguish BA and QA; and iii) the comparisons using only BA samples (that generated very low descriptor values) serves as a baseline that indicates that the partial separation that we obtain between BA and QA reflects some general differences between these two types of muscles.

We have analyzed the most frequent characteristics in **Table 2** to try to understand which is based on the difference in the organization between QA and BA. Interestingly, the six more frequent characteristics of the 1,000 BA-QA comparisons (all appearing in more than the 25% of the cases) are features that also are highlighted in the slow/fast cell randomization assay. This means that the feature selection method is considering the characteristics that capture the slow/fast mosaic as the most relevant to distinguish BA and QA organization. The most frequent characteristic is the “S. D. Area of slow cells” indicating the high relevance of the homogeneity in sizes for the slow cells in BA in contrast to the wider range of sizes in the case of QA **(Table S3).** The second (appearing in almost half of the cases) is “slow Neighbours of fast cells”. This characteristic would reflect the combination of the difference in the percentage of the slow cells between BA and QA together with the particular arrangement of slow and fast cells in the BA tissue. The third and fourth characteristics are the Average Strengths of fast and slow cells respectively. These two characteristics combine the information about the size and the number of neighbours of each type of cells. This appears slightly more relevant that the “average area of fast cells” (the fifth characteristic) to distinguish between both types of muscles. To finish with the more frequent characteristics we find “S.D. Neighbours of slow cells” that again reflect the fact that slow cells in BA are less variable due to their more constant size. All these characteristics are used by the method to separate both types of samples in a majority of combinations, therefore, in spite of the large heterogeneity between samples, we are able to conclude that the distribution of the two cell types is relevant to differentiate both sets of images.

### Biceps brachii present a distinct organization derived from the smaller size of the slow fibres with respect the fast fibres

We have explored the possible influence of the distribution of the slow and fast fibres in the global organization of BA and QA tissues. First, we have evaluated the importance of the percentage of each type of fibre (Δ proportion) in the organization of the tissue. We observe that driving this characteristic to a limit by choosing two sets of images from BA with a very diverse Δ proportion, we are able to obtain a clear separation in the PCA graph **(Fig. 3A).** However, the values for the PCA descriptor in the 1,000 combinations of BA-BA and QA-BA are clearly lower and do not correlate with Δ the “proportion” **(Fig. 3C, D).** We are convinced that this latter result is biologically relevant. It is clear that an abnormally high value for the Δ “proportion” of both sets is going to affect all the characteristics analysed. Nevertheless, the 1,000 combinations reflect a lot better the heterogeneity that can be found in the normal muscles among different individuals. Our analysis of the Δ “proportion” toss another interesting result, a comparison with very low Δ “proportion” can still present some differences as in the case shown in **Figure 3B.** This strongly support that other factors apart from the percentage of fibres are playing a role in the organization of the muscle tissue. To identify these factors we have analysed the muscle images as an arrangement of convex polygons. In these natural tessellations the area of the cells and their polygon sides are related in a way that affect the whole organization of the tissue (Sanchez-Gutierrez et al., 2016). The distribution of slow and fast fibres areas is slightly different in the case of BA **(Fig. 4B).** We interpret that the reduced size of a large part of the slow cells in BA affects the polygon distribution of each type of fibre. This is supported by the significant difference in polygon distribution between the slow and fast fibres in biceps brachii, and the increment of heptagons and octagons in the subpopulation of fast fibres **(Fig. 4D).** In the other hand, QA does not present significant differences between slow and fast fibres polygon distributions. This suggests that in the QA case that the type of fibre does not bias the organization of quadriceps. To confirm this hypothesis and deeper investigate the existence of organizational differences between QA and BA we have used a computational simulation **(Table 3** and **Figure 5).** For each image, we obtained 10,000 variations where the distribution of the slow and fast fibres was random. In this way we have been able to compare the real values for the 34 characteristics that are related with the distribution of the type of fibres **(Table 1**) with 10,000 random values. We think that this is a very robust baseline to compare with. We assume that if there is not an inherent organization of slow and fast fibres in the real tissue, the randomization should not affect these values. This is the case for QA, where only two of the 34 characteristics present a value out of the range obtained with the 10,000 randomizations **(Table 3).** On the contrary, in the BA experiment, almost half of the characteristics dramatically changed when compared with the real values. We conclude that there is a particular intrinsic arrangement in BA, and that the randomization largely alters this predetermined order. The analysis of the features that deviates from random, together with the integration of the whole set of data extracted, point out what are the basis of the biceps brachii organization. We propose that the difference in the size of slow and fast fibres impose the observed differential polygon distribution between both types of cells. In a packed tissue, small cells (slow) have less number of sides and associate with bigger cells (fast) with at higher frequency (due to Lewis law (Lewis, 1928). Following this argument, for example, a characteristic such “fast neighbours of slow cells” should have a bigger value than the random distribution. This is the case **(Table 3).** The analysis of the characteristics that are different in BA with respect the randomization provides us information to establish a model of how fibres organize in BA muscle **(Fig. 6A).** We propose that there is a tendency to the apparition of isolated slow fibres (small with low number of neighbours) in biceps brachii. This will affect the whole organization inducing homogeneity in the distribution of both types of fibres. As a result, there will not be large regions occupied only by fast fibres. This would be different to what happen in the “schematic” QA muscle **(Fig. 6B),** where some slow fibres are isolated and others are grouped without any tendency governing this organization. For this reason, the randomization assay generate values for most of the characteristics in the same range that the real QA values.

**Figure 6.**
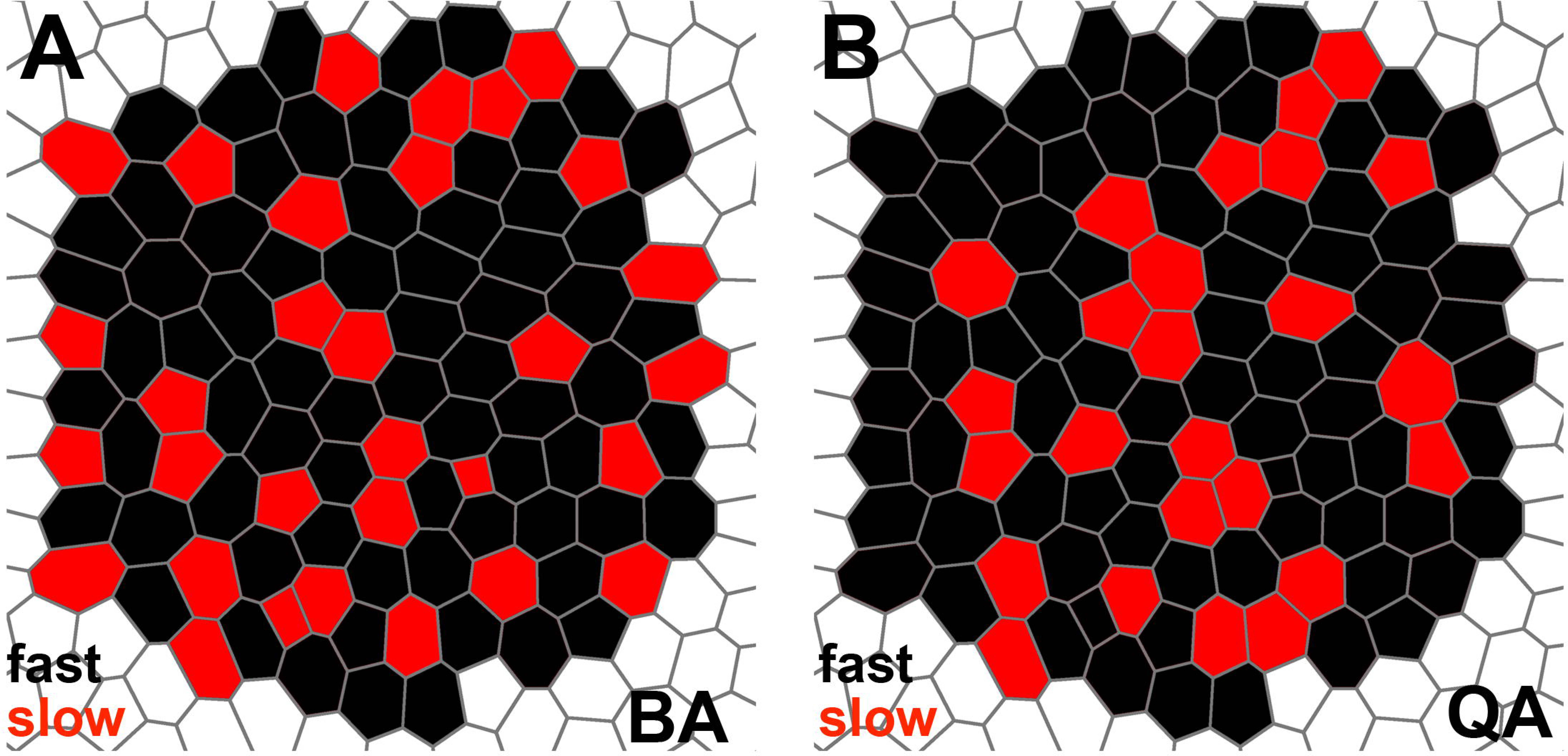
Scheme reflecting the different organization of BA and QA. Slow fibres are labelled in red and fast fibres in black. A) In the BA a high tendency for slow cells to be isolated govern the organization of the tissue. This induces a homogenous distribution of both types of fibres. **B)** There is no clear tendency in the organization. Slow fibres can appear isolated or grouped. The distribution is random.

## Conclusions

In this work we describe an organizational characteristic pattern based on the differential size of two different types of cells. Although a high heterogeneity exists among the analyzed samples, we have been able to detect the signature that generally distinguishes the biceps brachii from the quadriceps muscles. This is based on their slow/fast fibre organization. Our results clearly indicate that the relative bigger size of fast fibres in the biceps brachii is the origin of the intrinsic order that place homogenously the slow fibres on the tissue. In the other hand, quadriceps does not present any bias in the arrangement of both types of fibres.

These results are relevant from a translational point of view. A wide range of pathogenic changes have been described in the skeletal muscle of patients suffering from different neuromuscular diseases, both neurogenic and myopathic disorders. Subtle differences in the way of response to a pathogenic condition from one muscle to another could improve the diagnosis in early stages of the disease, which is the goal for any therapeutic intervention in this group of disorders (Kohn et al., 2014; Laing et al., 2011; Raman et al., 2015). Our results contribute to identify early changes associated to the fibre type distribution in the pathologic muscles, which would improve the early diagnosis and, therefore, would allow the application of a potential therapy before the muscle start a significant degeneration.

Muscles are not the only packed tissues where more than one cell-type can be found. During morphogenesis epithelial cells differentiate into precursors that are maintained within the epithelium during some time. This is the case of the neural crest of vertebrates (Duband, 2006) or the *Drosophila* sensory organs mother cells (Cohen et al., 2010). Still more intricate are some adult tissues in homeostasis, such the *Drosophila* midgut where enteroblasts, stem cells, enterocytes and enteroendocrine cells are integrated in the same layer (Lemaitre and Miguel-Aliaga, 2013). In all these examples the relative organization of the different cells types can be relevant for their function. Here we have described a new framework to analyze complex packed tissues where epithelial cells start to differentiate and more that one cell type is founded.

## Materials and Methods

### Tissue sampling

For the retrospective analysis of control male muscle tissue, we obtained images from processed biopsies stored in tissue banks at the Virgen del Rocío University Hospital (Seville). Our database consists of 90 ROI extracted from 34 images which were selected from 14 biopsies for biceps brachii Adult (BA) and 25 ROI extracted from 9 images which were selected from 6 biopsies for quadriceps adult (QA). We selected a ROI with resolution 1,000x1,000 pixels from images of 3,072x4,080 pixels. In this way it is possible to avoid small artefacts due to the manipulation and staining of the samples. All biopsies were performed under informed consent using a standardized protocol (Dubowitz and Sewry, 2007) and ROIs were processed as described in (Sáez et al., 2013b). In this way, a fibre segmentation, in which fibre contours were detected, was performed in order to extract information relative to fibre and collagen content.

### Geometric and network feature extraction

Geometric features such as the fibre area or the length of the major and minor axes of the fibre can be extracted from the detected contours. A network of fibre to fibre contacts was derived form the segmented image following the steps described in (Sáez et al., 2013b). This allowed to obtain other parameters that take into account the neighbouring vicinity of each fibre, such as the ratio between the fibre area and adjacent fibre areas, or the ratio between the fibre area and the area resulting from the expansion of its contour (computed in the previous step). Finally, features extracted from graph theory applied to the muscle network were also computed (values for all characteristics in each image in **Table S1**).

In this work a total of 69 characteristics have been computed. They included 14 geometric features, 20 features derived from the muscle network, 34 from graph theory and 1 last characteristic which gave us the proportion of slow cells **(Table 1).** We defined 3 subsets of characteristics in order to employ it in different comparisons. First set was performed by all 69 characteristic computed. Second set was defined by 35 characteristics related with slow and fast cell information (in bold in **Table 1**). Third set was composed exclusively by 14 geometric characteristics (14 first features in Table 1) and the proportion of slow cells.

### Principal Component Analysis features selection

A feature selection step was performed to analyze the discrimination power of a set of characteristics of one’s mentioned above that distinguish better two groups of images. The method selects and evaluates features using Principal Component Analysis (PCA) and PCA’s descriptor that quantify the degree of separation between the two groups of images that are compared (Sanchez-Gutierrez et al., 2013). We have tested every possible combination of three features in the first iteration and applied the PCA. The method keeps the ten combinations of three features with higher PCA descriptor value. In the second iteration, all features are individually tested again in combination with the ten trios of features. Again, all the combinations are evaluated and the program keeps the five with higher PCA descriptor value for each one of the ten trios. Therefore, at this step the program handles 50 quartets of features. In the next iteration, the same process is repeated but only two best features are added, accumulating 100 quintets of features. The process continues adding only one feature per iteration step. The iteration process is stopped when seven features have been selected or when the value for the PCA descriptor is lower than in the previous step. Finally, we chose the ensemble of features that presented the highest value for the PCA descriptor among the 100 groups.

### Comparison of BA and QA images

Due to the large difference in the number of ROIs (90 BA vs. 25 QA) we designed a protocol to use all the available samples and at the same time be able to obtain comparable results. We employed a random process of sample selection to be able to compare the same number of samples each time. We selected “25” random ROIs (the smallest quantity of ROIs in one of the groups) to perform the PCA features selection described above. To be sure that we used all the available samples we did this process 1,000 times to perform 1,000 comparisons. Therefore, for each comparison, we also obtained 1,000 PCA descriptors and 1,000 sets of relevant characteristics. In order to know which characteristics were most relevant to discriminate two categories along using all the available images, we calculated the rate of appearance of each feature between the selected ones. Table 2 and Table S3 show the most frequent characteristic in each comparison performed in this study.

### Relation between discrimination power and slow fibres proportion

To test if there is a correlation between the values of the PCA descriptors obtained with the 1,000 comparisons and their proportion of fast and slow fibres, we defined the value Δ “proportion” per each one of these 1,000 comparisons. “Δ proportion” was calculated as the difference between the average percentages of slow cells between two groups analyzed in each one of 1,000 comparisons. The Pearson’s correlation coefficient was obtained to analyse the possible correlation between the value of the PCA descriptor and the slow fibre proportion.

### Statistical differences between BA and QA fibre characteristics

We used Multivariate Analysis of Variance (MANOVA) test to perform three comparisons of the polygonal distributions: a) BA total fibres vs QA total fibres, b) BA fast fibres vs BA slow fibres, c) QA fast fibres vs QA slow fibres **(Table S4).** If *p-value* <0.05, distributions were considered to be significantly different. The MANOVA tests were performed using only the values for cells with 4, 5, 6, 7 and 8 sides. We discarded the cells with 3, 9 and 10 sides, since they were not present in all the images. In the three comparisons above we also analyzed the differences between the values for each type of polygon. First, we evaluated if the two compared categories values presented similar distribution and variance using Kolmogorov-Smirnov and F-Snedercor tests respectively. In case that data presented different distribution and a different variance, we employed Wilcoxon test to compare the means from both groups. We employed a two tails Student’s t-test to compare the means in the cases where both distribution and variance of the two sets of data were similar **(Table S4).**

We used the two samples Kolmogorov-Smirnov test to compare “log10 Normalized Area” distribution of each category of “BA fast fibres vs BA slow fibres” and “QA fast fibres vs QA slow fibres”.

### Slow and fast cell randomization

In order to know how the spatial distribution of slow and fast cells affected to the organization of the muscle, we randomized the positions of fast and slow cells without altering their proportion. In each ROI, every cell was labelled as “fast” or “slow” randomly, maintaining the relative number of fast and slow cells. This process changed the values for the 34 characteristics related with fast and slow properties. We performed 10,000 randomizations for each ROI. For each category and randomization we calculated the average value of each one of the 34 characteristics. To obtain the “original” value for each characteristic we averaged the values of all the available images (90 for BA and 25 for QA). We plotted the distribution of 10,000 values for each characteristic and compared its minimum, maximum, and median values with the “original” average value of slow and fast cells. **(Fig. 5, Table 3**)

### Polygon and area distribution calculations

We analyzed polygon and area distribution in our images to investigate about the organization of fast and slow cells in relation with their size **(Fig. 4).** To make the polygon distribution graphs with the corresponding error bars for each category (BA, BA slow cells, BA fast cells, QA, QA slow cells and QA fast cells) cells were grouped by biopsy.

To compare Area from different categories, we calculated the Normalized Area:

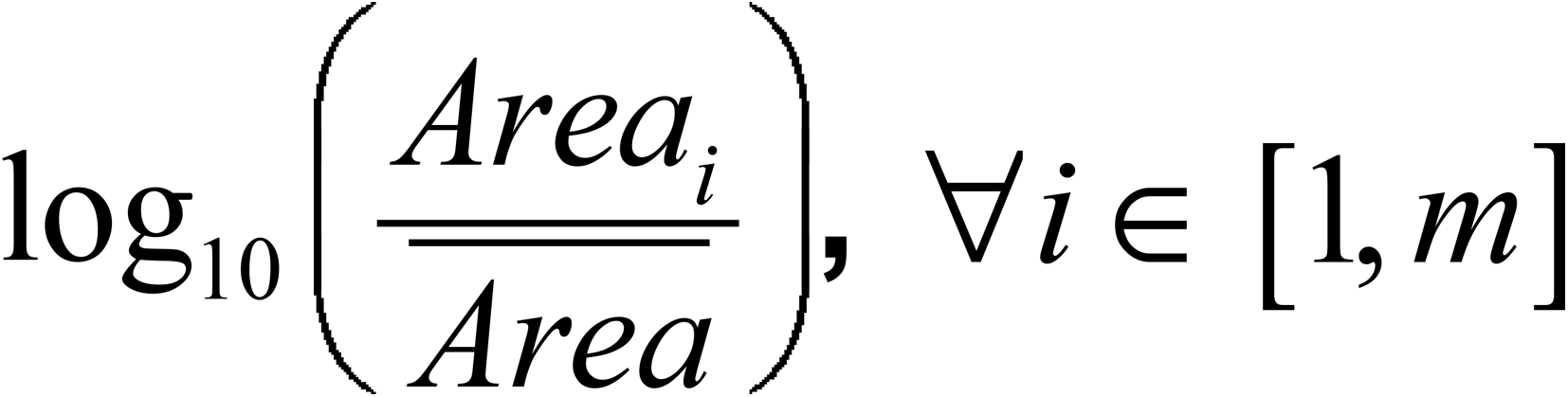

Where m is the number of cells in the image, *Area_i_* is the size of the cell and 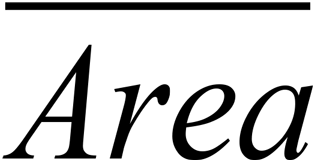is the mean Area of all the valid cells. We classified the values in bins of 0.02 units to visualize the Normalized Area distribution. The use of the log10 makes the values distribute similar to a normal distribution facilitating the comparison.

